# Tracing the Indian Population Ancestry by *cis*-linked Mutations in *HBB* Gene

**DOI:** 10.1101/2021.04.18.440318

**Authors:** Amrita Panja, Prosanto Chowdhury, Anupam basu

**Affiliations:** Molecular Biology and Human Genetics Laboratory, Department of Zoology, The University of Burdwan; Institute of Child Health, Kolkata

**Keywords:** *HBB* gene, mutation, migration, thalassemia, Indian population

## Abstract

**Background:** Human left their genetic footprints during the time of migration throughout the different countries all over the world. Human evolution was studied through various markers. India is a country of rich heritage and cultural diversity. The modern Indian population is derived from two ancestral groups, viz.-Ancestral North Indians (ANI) and Ancestral South Indians (ASI).

**Aim:** Finding out the migratory route of the modern Indian population by studying ‘cis’ acting mutations of human beta-globin (*HBB*) genes.

**Subjects and methods:** A total of 120 thalassemia subjects were enrolled. DNA sequencing was done for mutation detection in the *HBB* gene. Some previous literature reviews were gone through for tracing mutations, all over the world and in the Indian subcontinent.

**Results:** Nine thalassemia patients were found where *HBB*:c.92G>C and *HBB*:c.-92C>G mutations co-exist together in ‘*cis*’ condition. Only one patient had *HBB*:c.51delC and *HBB*:c.33C>A. The pedigree analysis confirmed the presence of these mutations in ‘*cis*’ condition and vertical transmission from one generation to the next. Literature reviews also reassure the co-existence of these mutations from different countries.

**Conclusion:** The co-existence of these ‘*cis*’ acting mutations helps to point out the possible migratory route of ANI population after venturing out of Africa.

## INTRODUCTION

Genetic evidence suggests that the world’s most significant population divergence occurs among the African hunter-gatherers nearly 3000, 000 years before the present. With the help of DNA sequencing technology, full genome sequences of prehistoric ancestral Africans can be traced. The modern genetic diversity of Africans occurs due to migration intended for pastoralism and agriculture in the last few thousand years (Schlebusch and Jakobsson, 2018). Geographical migration indicates the movement of people from one dwelling to another, although it is a heterogeneous and complex process (Mascie-Taylor and Krzyżanowska, 2017). In the previous three decades, several genetic studies have been conducted to explore the human evolutionary history of migration. So far, more than a few markers have been used for tracing the ancestry of human civilization. The African origin of anatomically modern human (AMH) is reinforced by genetic, paleoanthropological and archaeological studies (Stringer C, 2002; White et al., 2003; Ramachandran et al., 2005; Jakobsson et al., 2008). Earlier, protective allelic variants of the few genes, including Duffy antigen/chemokine receptor (*DARC*), Glucose-6-phosphate dehydrogenase (G6PD), Lactase (*LCT*), Minichromosome Maintenance Complex Component 6 (*MCM* 6), Amylase (*AMY*1) and pattern of haplotypes have been extensively studied among the population of Africa for better understanding the migratory pathway in course of evolution (Jones et al., 2013; Priehodova et al., 2014; Fan et al., 2016; Fernandez et al., 2017). The distribution pattern of different mutations is population specific. Germline mutations act as the raw material for evolution and these are responsible for causing different inherited diseases. When the groups of ancestral human distributed in different countries all over the world; each population often avoided or altered mutations from the genome for better adjusting to the surrounding environment (Harris and Pritchard, 2017). It is assumed that specific Single Nucleotide Polymorphism (SNPs) account for approximately 15% whereas common ones stand for the rest of 85% (Spichenok et al., 2011; Nelson et al., 2012).

Reich and his colleagues studied on 25 different ethnic groups and based on the findings of mt-DNA and Y-chromosome, they concluded that the modern Indian population is derived from basically two ancestral groups, viz.-Ancestral North Indians (ANI) and Ancestral South Indians (ASI) (Reich et al., 2009). The ANI people have genetic similarities and affinities with the Middle East, Central Asia and Europe. On other hand, the ASI is related to indigenous Andaman Islanders. Due to multiple rituals, traditions, widely variable social customs and interracial marriage system, random genetic admixture was observed among Indo-European–speaking upper castes. Therefore, the present Indian population is the admixture of ANI and ASI (Reich et al., 2009; Moorjani et al., 2013). On the other way, it has been demonstrated by DNA microarray-based autosomal SNP analysis, that there are additional ancestral groups in India apart from the previously mentioned (Basu et al. 2016). Five ancestral groups are residing in the country, including Indo-European (people of Northern, North-Western, Eastern, Western part of India and mainly belongs to upper caste in a social hierarchy including Khatri, Gujarati Brahmin, West Bengal Brahmin, Maratha), Dravidian (Southern part and mainly belongs to Lower-middle caste and tribal communities including Iyer, Pallan, Kadar, Paniya), Austro-Asiatic (inhabitants of the Central and Eastern part of the country and belongs to tribal ethnicity including Ho, Santal), Tibeto-Burman (found in North-Eastern part of the country and mainly from upper caste including Manipuri Brahmin, Tripuri, Jamatia), Indo–European (residents of Northern India and belongs to the tribal community) and Ongan (tribal communities including Onge and Jarawa in Andaman and Nicobar island) (Basu et al., 2016; Mahal and Matsoukas, 2018).

Human beta-globin gene (*HBB*) is situated on chromosome 11 and it provides information for beta-globin chain synthesis of a larger protein called hemoglobin. Mutations in *HBB* gene causes hemoglobinopathies including thalassemia. There are more than 500 mutations so far documented in HbVar Database (Giardine et al., 2014). A population-based study from different regions of the Indian subcontinent (Gujarat, Punjab, Sindh and North-Western states) revealed the time of origin of different thalassemia mutations (Gadhia et al., 2019; Shah et al., 2017). Moreover, there are several single nucleotide polymorphisms (SNPs) which co-exist with thalassemia mutations. So far, few haplotype-based studies have been conducted among the thalassemia patients of India (Gupta et al., 2008; Nongbri et al., 2017).

In this present investigation, it has been hypothesized that the *cis*-linked β-thalassemia mutation pair can act as the genetic marker for exploring the ancestry of the Indian population. Accordingly, we have investigated the linked thalassemia mutation pair present in the studied Bengali thalassemia subjects. Moreover, we have gone through the pieces of literature for finding out the existence of these observed *cis*-linked β-thalassemia mutation pair in other populations throughout the world to establish the haplotype markers in finding out the Indian population ancestry and this will be the first of its kind for this population.

## MATERIALS AND METHODS

### Subject information

The study was approved by the Institutional Ethic Committee of The University of Burdwan and conducted as per Helsinki declarations. The Information sheet was provided and written consent was taken from each patient included in this study. A detailed description of the purpose of the study was disclosed to the participants. All the experiments during this study were accomplished following relevant guideline and proper regulations. Samples were collected from Burdwan Medical College and Hospital. Subjects included in this study came from different districts across West Bengal and they mostly belong to Bengali ethnicity.

### Inclusion criteria

The subjects positive for thalassemia screening tests, including HPLC (Hemoglobin A2 i.e HbA2 of >4.0% and/or fetal hemoglobin i.e. HbF of >2.0%) and regular blood tests (mean corpuscular hemoglobin i.e. MCH of <27pg, mean corpuscular volume i.e. MCV of <80fl, hemoglobin <9g/dl) were included in this study. Clinical and hematological data were collected from the medical records.

### Exclusion criteria

The subjects with iron deficiency anemia or other clinical complications were excluded from this study. Any unconfirmed blood transfusion-dependent patients were excluded from this study.

### Subject size

In the present study, 120 thalassemia major subjects were included.

### Sample collection

About 2-3 ml of blood samples were collected in EDTA vacutainers following standard phlebotomy procedure. Sampling was done in an interval of at least 4 weeks from the last transfusion date or just before the next transfusion. Blood samples were stored at -80 °C until DNA extraction.

### Genomic DNA extraction

Genomic DNA was extracted from peripheral blood lymphocytes using the standard phenol-chloroform extraction method with some modifications (Sambrook and Russell, 2006).

### Human beta-globin (*HBB*) genotyping by DNA sequencing PCR

PCR was carried out to amplify the part of *HBB* gene, including 5’UTR, Exon 1, Intron 1and part of Exon 2. Briefly, the amplification reaction included 35 cycles of denaturation at 94° C for 1 minute, annealing at 45 °C for 50 seconds, extension at 72° C for 50 seconds. 1% agarose gel electrophoresis was done to extract the 800bp amplified PCR product. The purified PCR products were subjected to direct sequencing with an ABI 3100 DNA Sequencer (Applied Biosystems; Foster City, CA, USA).

### Selection of ‘*cis’* linked *HBB* mutation pair

In the studied group, two ‘*cis*’ linked mutation pairs of the *HBB* gene were selected.These are (i) *HBB*:c.92G>C and *HBB*:c.-92C>G (ii) *HBB*:c.51delC and *HBB*:c.33C>A. We obtained only one proband with *HBB*:c.51delC and *HBB*:c.33C>A mutations and nine with *HBB*:c.92G>C and *HBB*:c.-92C>G mutations in the ‘*cis*’ condition. Thus, we studied the pedigree for at least three generations for the proband having *HBB*:c.51delC and *HBB*:c.33C>A. mutations. On the other hand, in case of *HBB*:c.92G>C and *HBB*:c.-92C>G mutation pair, the pedigree of each affected family was analysed for at least two generations.

### Literature reviews on the occurrence of *HBB*:c.51delC and *HBB*:c.33C>A & *HBB*:c.92G>C and *HBB*:c.-92C>G mutations

A comprehensive HbVar database (Giardine et al., 2014) searching was done for finding out the occurrence of this paired mutations in world populations. For better understanding, we have categorized the prevalence of these mutations continent wise throughout the world. At the same time, the occurence of these studied mutation pairs in other states of India was also thoroughly revised. The origin and distributions of the susceptible communities were studied by going through the literature reviews. Here in the present study, we have divided the geographic region of India into five zones, Zone-1: North India (Jammu and Kashmir, Uttar Pradesh. Punjab, Haryana), Zone-2: Central India (Madhya Pradesh, Chattisgarh), Zone-3: West India (Gujarat, Maharashtra, Rajasthan), Zone-4: South India (Kerala, Karnataka, Andhra Pradesh), Zone-5: East India (West Bengal, Assam, Orissa, Bihar, Jharkhand). The occurrence and distribution of these *cis*-acting mutation pair were described according to different zones in the country.

### Data Analysis

All the data were entered into and managed using Microsoft Excel 2007 (Microsoft, Redmond, WA, USA). The gene mutations, frequency, distribution and spectrum of thalassemia were analysed precisely with a descriptive method. Studied subjects carrying the mutations were collected with their medical data and laboratory parameters. The patients carrying the associated mutations were further distinguished by their ancestor origin. The Pedigree analysis helped to reveal the ancestry of the forerunners of the affected families with the specific haplotypes.

The chromatograms obtained after automated DNA sequencing were interpreted by using Chromas Lite software (v 2.1). Moreover, DNA Baser (v 4.36.0.2) was used for DNA sequence assembly.

## RESULTS

### Co-existence of *cis*-linked *HBB* mutations among the studied thalassemia subjects

#### 1. *HBB*:c.51delC and *HBB*:c.33C>A

A 15-year-old girl came to our laboratory for the investigation of the possibility of thalassemia. She originated from the Birbhum district of West Bengal. She presented clinical features similar to thalassemia. DNA sequencing revealed the presence of both *HBB*:c.51delC and *HBB*:c.33C>A mutations along with *HBB*:c.79G>A mutation (Figure1). The pedigree analysis was carried out for three generations and it was found that *HBB*:c.51delC and *HBB*:c.33C>A mutations were vertically transmitted from mother (II-8) to the proband (III-9). Consequently, the maternal aunt (II-4) of the proband had both *HBB*:c.51delC and *HBB*:c.33C>A mutations (Figure 2).

**Fig 1.**
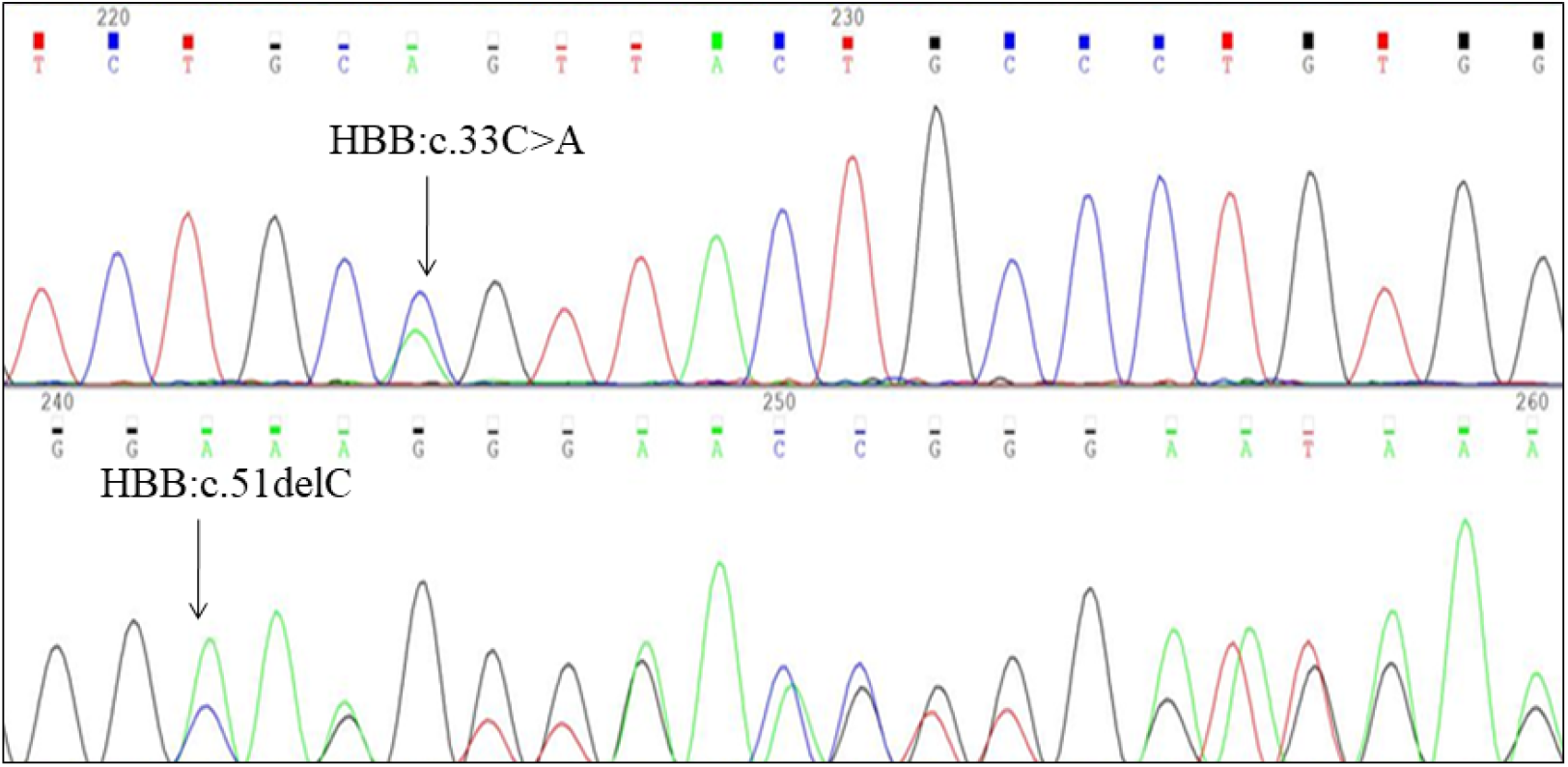
DNA sequence chromatogram of the selected region of *HBB* gene showing the presence of the substitution mutation *HBB*:c.33C>A, and *HBB*:c.51delC deletion mutation in ‘*cis*’-linked condition.

**Fig 2.**
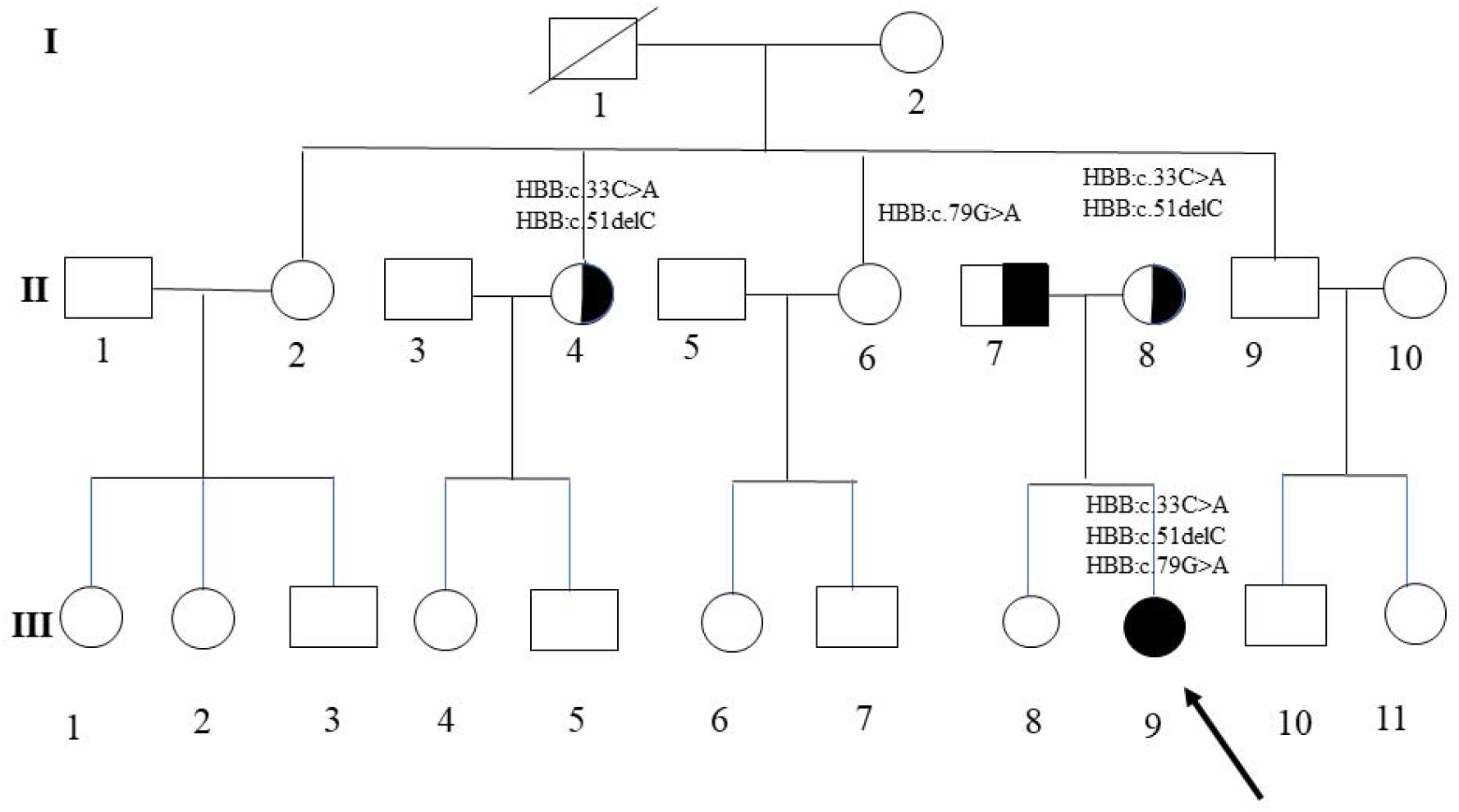
Pedigree of the family of proband where *HBB*:c.33C>A and *HBB*:c.51delC mutations were found to be co-existed together. The proband (III-9) got these paired mutations from her mother (II-8). II-4 had both *HBB*:c.33C>A and *HBB*:c.51delC mutations together.

#### 2. *HBB*:c.92G>C and *HBB*:c.-92C>G

DNA sequence analysis revealed that both *HBB*:c.92G>C and *HBB*:c.-92C>G mutations are inherited as cis -linked (Figure 3) haplotype. Pedigree analysis shows the transmission of these paired mutations from parents to their offspring irrespective of sexual biases.

**Fig 3.**
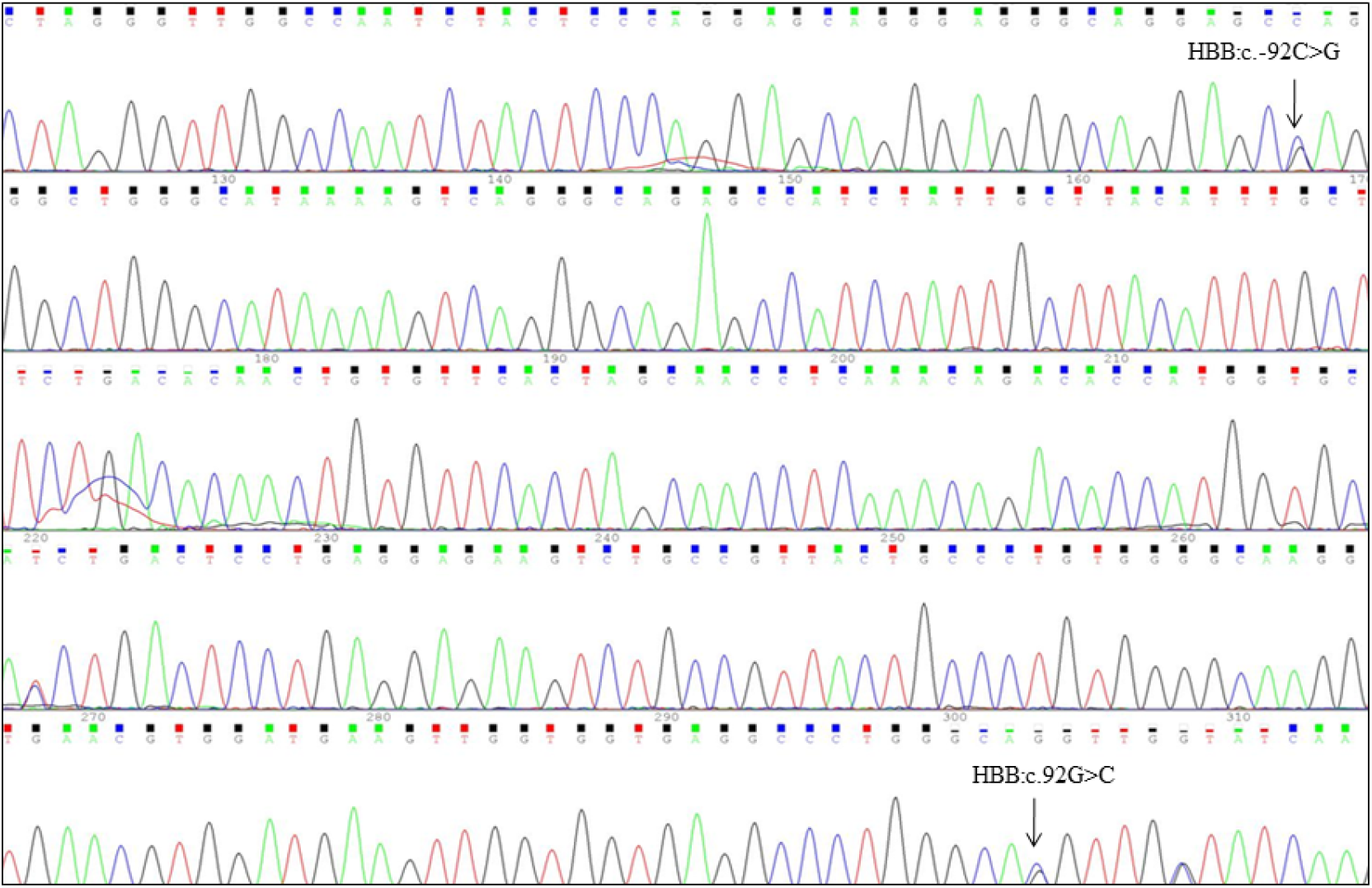
DNA sequence chromatogram of the selected region of *HBB* gene showing the presence of the substitution mutation, *HBB*:c.92G>C and *HBB*:c.-92C>G in ‘*cis*’-linked condition.

### Findings from literature reviews

#### 1. Haplotype distribution all over the World

Previously, it had been reported that *HBB*:c.33C>A mutation was a very rare polymorphism, which was an ancestrally originated allele, upon which *HBB*:c.51delC mutation exists (Old et al., 2001; Fisher et al., 2003). The co-existence of *HBB*:c.51delC and *HBB*:c.33C>A mutations were reported earlier from Saudi Arabia which is situated at the South-West part of Asian subcontinent (Abuzenadah et al., 2011). According to HbVar Database, this mutation pair was found previously from Azerbaijan (1.5%), Pakistani (2.2%), Pathan (3.81%), Punjabi (1.6%), Singapore (0.75%) population (Patrinos et al., 2004; Thein SL, 2013). This mutation was also found in Malaysia (Kadazandusuns in Sabah province, Penang and Kedah province) and Chinese ethnicity (Tan et al., 2010; Hassan et al., 2013; Tan et al., 2015; Islam et al., 2018). The population-based studies done from Sri Lanka, Thailand and Syria revealed the coexistence of both *HBB*:c.51delC and *HBB*:c.33C>A mutation in homozygous mutant condition (Old et al., 2001). Therefore, the haplotype has prevailed in the Eastern part of Africa, Eastern Europe, South-Western as well as South-Eastern part of Asian subcontinents.The worldwide distribution of this haplotype has been depicted in Table 1.

**Table 1.**
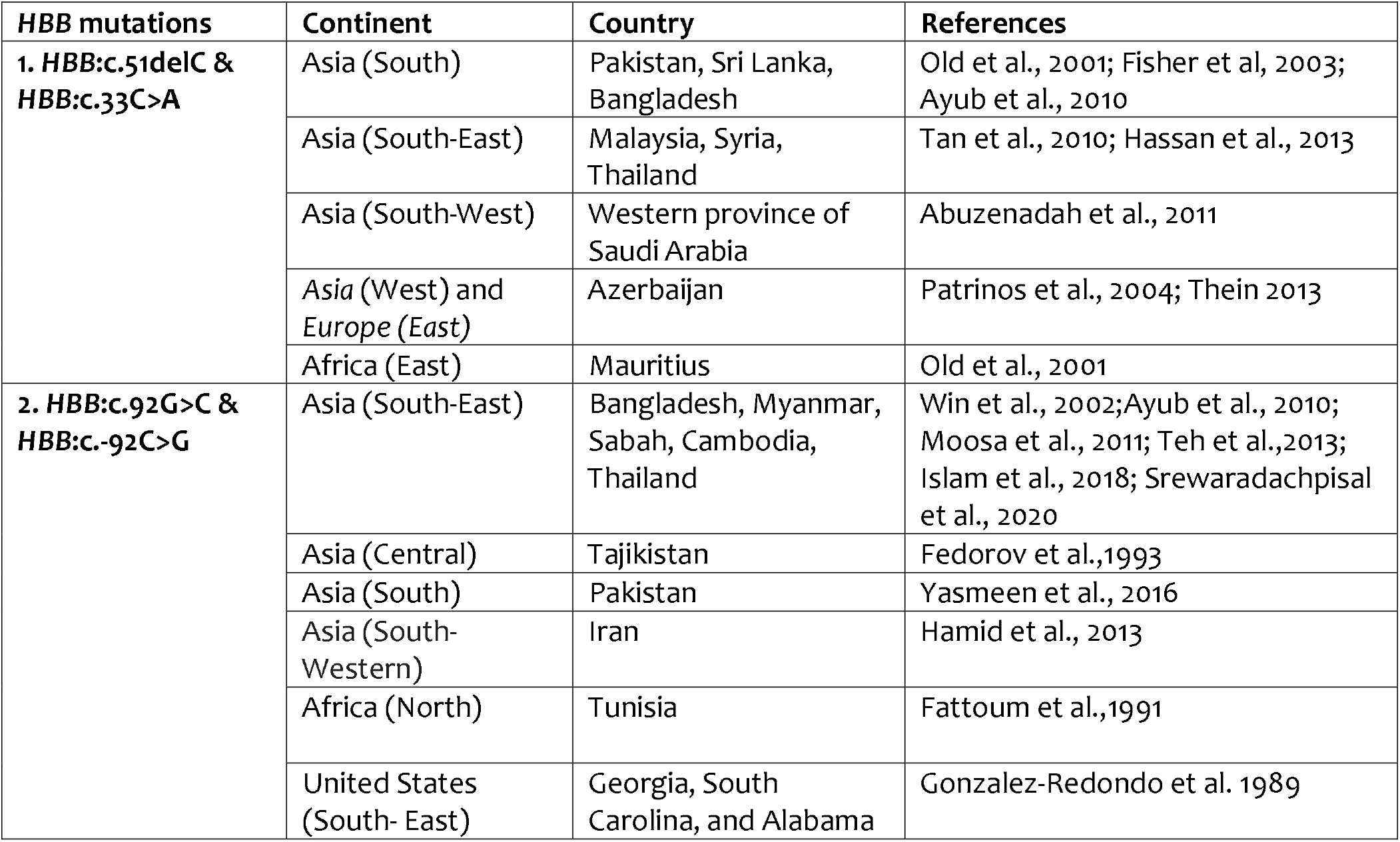
The list of countries with the corresponding continents where the population possessing *HBB*:c.51delC & *HBB*:c.33C>A and *HBB*:c.92G>C & *HBB*:c.-92C>G mutations in *cis*-condition

*HBB*:c.92G>C (Hb Monroe) was first observed in a transfusion-dependent β-thalassemia major 15-year-old female of USA (Gonzalez-Redondo et al., 1989). The combination of Hb *HBB*:c.92G>C and *HBB*:c.-92C>G mutation was reported earlier from South Asian countries like Tajikistan and Pakistan (Fedorov et al., 1993; Yasmeen et al., 2016). Five individuals having thalassemia minor phenotype from four unrelated families of Khuzestan of Southern Iran was found to have the mutation pair (Hamid et al., 2013). Thus, the haplotype mutation prevails in the countries situated on the South-western countries of Asian continent. Consequently reports are available from South-East Asian contries including Bangladesh, Myanmar and Thailand (Fattoum et al., 1991; Win et al., 2002; Ibn Ayub et al., 2010; Moosa et al., 2011; Teh et al., 2014; Islam et al., 2018; Srewaradachpisal et al., 2020) (Table 1).

#### 2. Haplotype distribution in India

In India, a community-wide study revealed the co-existence of *HBB*:c.51delC and *HBB*:c.33C>A mutations from North India (Zone-1) (2.1%-3.6%) (Gupta et al., 2003; Meena et al., 2013). One of the studies done among the population of Uttar Pradesh shows this mutation was mostly found in Backward classes (10-50%), moderately among Vaishyas, Kshatriyas communities and less common between Brahmins and Muslims (<10%) (Sinha et al., 2009). It was also reported from Central India (Zone-2) including Madhya Pradesh (Pawar et al., 1997). Likewise, this haplotype was observed from the Western part of the country (Zone-3) including Gujarat (Kachhiya Patel, Dhodia Patel, Modh Bania communities), Maharastra (Aurangabad, Khandesh, Saurashtra) (Colah et al., 2010; Bhukhanvala, 2013). Interestingly, it was predominantly found among Brahmins in comparison to Rajputs and Vaishya caste in Gujarat (Vaz et al., 2000). One demographic study was done on East Indian (Zone-4) population revealed the frequency of this mutation is 3.6% (Nagar et al., 2015). A large scale population wide study done from different states of India demonstrated the frequency of this *cis*-linked mutations is 7.4% in Kerala, 5.1% in Uttar Pradesh, 2.25% in Maharastra, 1.8% in Bihar, 1.55% in Sindh, 1.13% in West Bengal (Pawar et al., 1997). The occurrence of this mutation in the different parts of India has been described in Table 2.

**Table 2.**
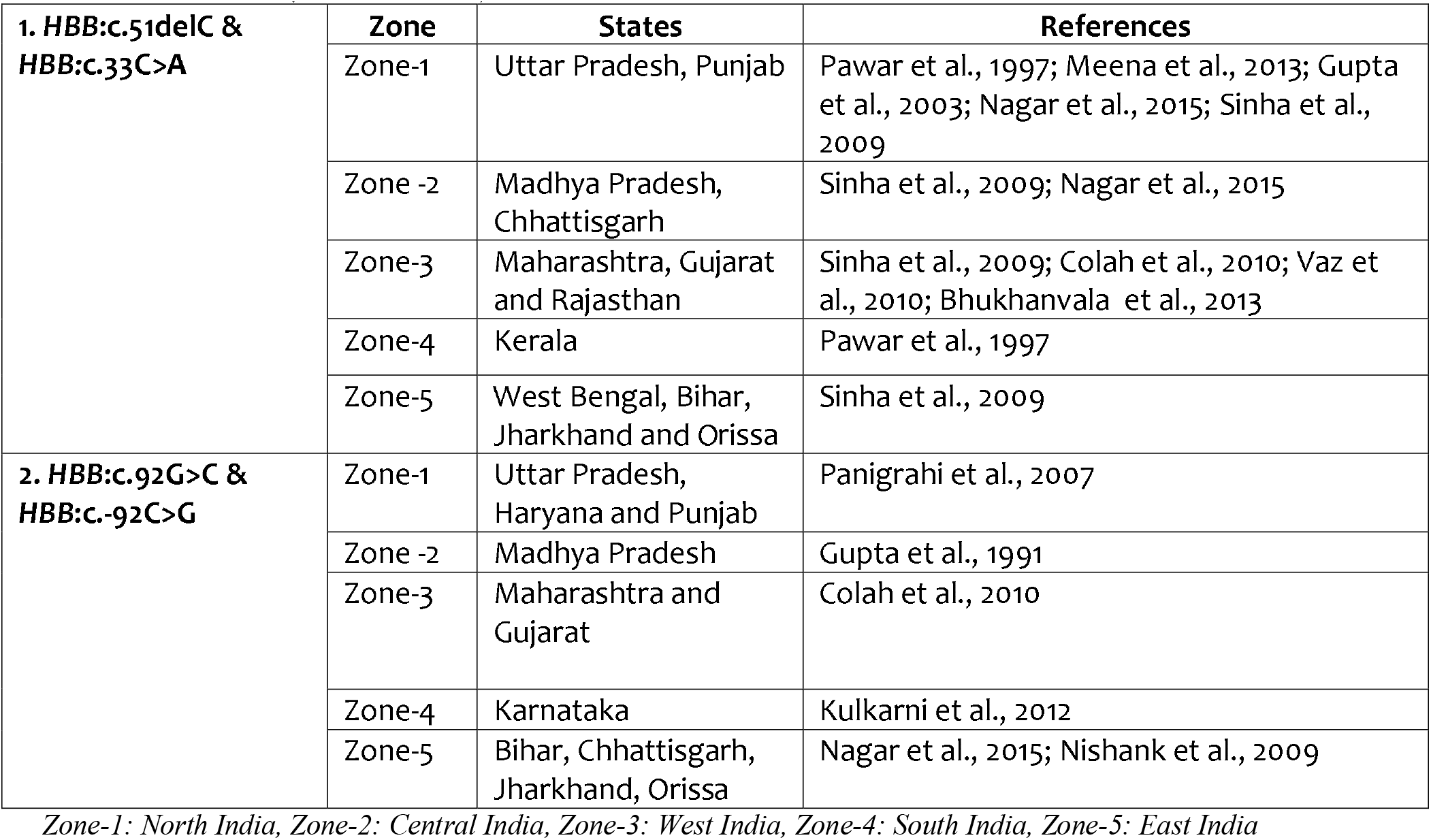
Zone wise regional distribution of *HBB*:c.51delC & *HBB*:c.33C>A and *HBB*:c.92G>C & *HBB*:c.-92C>G mutations (‘*cis*’-condition) in India

Hb Monroe (*HBB*:c.92G>C) was first detected among the Gond tribal community of Central India (Zone-2) where the interaction between α and β-globin chain variants has been demonstrated for influencing the disease severity (Gupta et al., 1991). It was earlier found from Northern part of India (Zone-1) including Uttar Pradesh, Haryana, Punjab; Western part (Zone-3) containing Gujarat, Maharastra and Eastern part (Zone-5) counting Orissa (Panigrahi et al., 2007; Nishank et al., 2009; Colah et al., 2010; Nagar et al., 2015; Nigam et al., 2017; Bakhle et al., 2018). A large scale demographic study done in different states of India revealed the overall frequency of this mutation in India is 2.6%. The prevalence rate is 2% in West India (Zone-3), 1.8% in North India (Zone-1), 2.7% in Central India (Zone-2), 1.7% from South India (Zone-4), 5.8% in East India (Zone-5) (Sinha et al., 2009, Kulkarni et al., 2012; (Table 2).

### Prediction of human migratory pathway in Indian subcontinent using *cis-*acting *HBB* mutation pairs

According to some previous reports, the co-existence of *HBB*:c.51delC, *HBB*:c.33C>A and *HBB*:c.92G>C, *HBB*:c.-92C>G mutations are mostly confined to Western Eurasian countries, including Saudi Arabian, Turkey, Iran, Azerbaijan and Pakistan. Moreover, these two haplotypes were also observed in South-East Asian countries including Bangladesh, Myanmar, Thailand and Malaysia (Fedorov et al., 1993; Abuzenadah et al., 2011). In India, majority of the earlier reports on these haplotypes are available from Northwestern (Zone-1 and 3), Central (Zone-2) and Eastern part (Zone-5) of India including Uttar Pradesh, Punjab, Haryana, Gujarat, Maharastra, Madhya Pradesh, Bihar, West Bengal (Sinha et al., 2009; Colah et al., 2010; Nagar et al., 2015). Interestingly, in the Southern part of India (Zone-4), the prevalence of these two studied *cis*-acting mutation pair is very scanty. Therefore, the existence of these mutations helps to highlight the migratory path of Indo-European as well as ANI population in India and neighbouring countries. The Southern states (Zone-4) of India (Andhra Pradesh, Karnataka, Tamil Nadu and Kerala) contain a predominantly Dravidian population, ethnically and culturally quite discrete from the largely Indo-European populations of Northern, Central and Western India (Reich et al. 2009).

## DISCUSSION

In the present study, we have found two genetic haplotype markers related to thalassemia; the first pair is comprised of *HBB*:c.51delC and *HBB*:c.33C>A and the second is of *HBB*:c.92G>C (Hb Monroe) and *HBB*:c.-92C>G. These ‘*cis*’-acting mutation pairs are closely linked or co-existed with each other from one generation to the next. Pedigree analysis of the affected families showed the vertical transmission of these two mutations from one of the parents to the proband.

Possibly, this study is the first pioneer effort where mutations of β-thalassemia help to enlighten the probable migratory tract of the modern human population, specially the ANI population of India. According to the findings of the present work, it can be assumed that after originating from Africa (nearly 200,000 and 60,000 years ago), a branch of the modern population migrated through the horn of North Africa, then entered into the Western part of Eurasia, greater Caucasian region, Saudi Arab, Azerbaijan (situated at the bank of the Caspian Sea), Turkey, Iran, Pakistan and then entered India through the North-Western boundary by invasion through Indus valley, Sindh province, Punjab and Gujarat (Figure 4). Moreover, the significant absence of these *cis*-acting *HBB* mutation in the Southern population of India may support the separate origin of the ASI and ANI group and the possibility of arriving by following the “southern exit” wave out of Africa (Basu et al., 2016). So, the co-association of thalassemia mutations helps to point out the possible migratory route of the ANI population and it supports the Indo–European lineage in the ancestral population of India (Figure 5). The findings obtained from the present investigation also reinforcing and supports the migratory path of anatomically modern human towards the Indian subcontinent same like that have guided by previous marker studies (Cordaux and Stoneking, 2003; Stanyon et al., 2009; Sharma et al., 2018).

**Fig 4.**
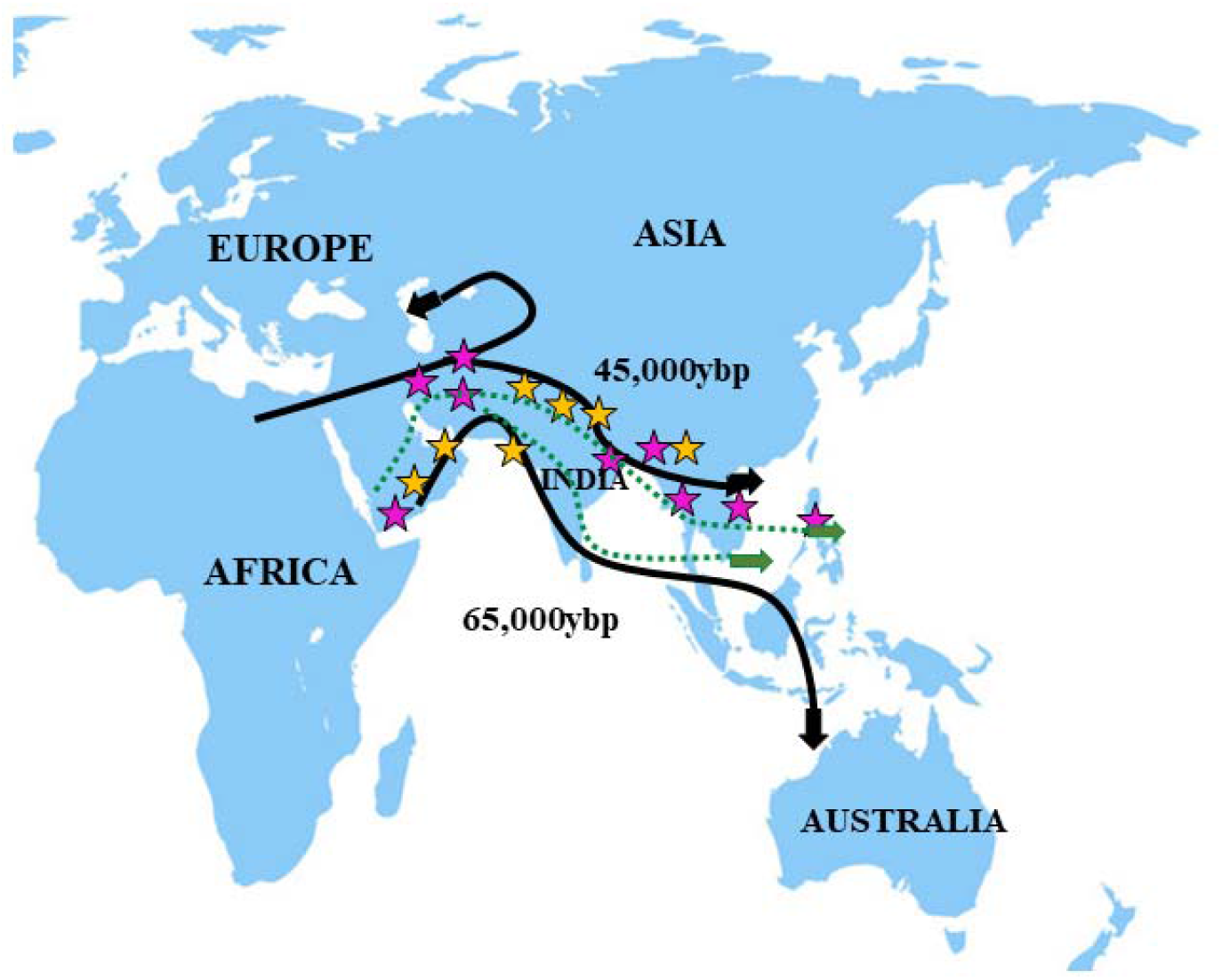
Mirroring of *HBB*:c.33C>A and *HBB*:c.51delC mutations [by yellow stars] and *HBB*:c.92G>C and *HBB*:c.-92C>G mutations [by pink stars] to the map of the routes of the human migration to the Indian subcontinent as defined by Tamang et al. 2012 [black solid line]. The green dotted lines which are drawn based on the position of co-associated mutations observed in the present study indicate the possible migratory route of modern human population after originating from Africa and it supports the Indo–European lineage in the ancestral population of India.

**Fig 5.**
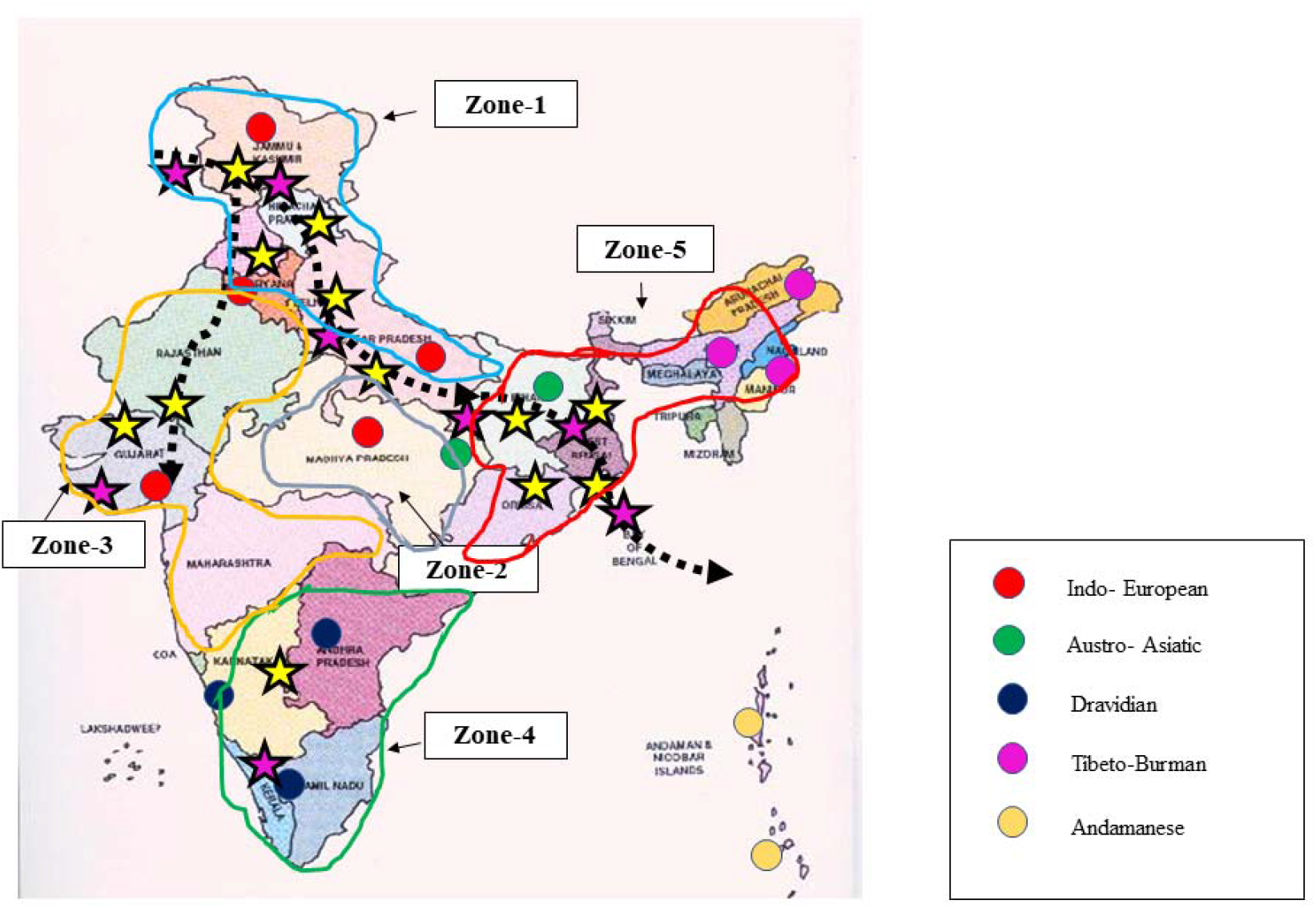
The co-existence of *HBB*:c.33C>A and *HBB*:c.51delC mutations indicated by yellow stars and *HBB*:c.92G>C and *HBB*:c.-92C>G mutations by pink stars to the population ancestry map of India designated by Reich et al. 2009. The black coloured broken lines indicates the probable migratory route of modern Indian population as well as lineage of Anestral North Indians (ANI) based on co-associated mutation pair, *HBB*:c.33C>A, *HBB*:c.51delC and *HBB*:c.92G>C and *HBB*:c.-92C>G. The prevalence of these ‘*cis*’-linked mutation pair has been described under five zones (Zone1 to 5)

For genetic diversity study, uniparental and biparental markers have been investigated for the last few decades. Among the studied markers, Y-haplogroup, mitochondrial DNA (mt-DNA), microsatellite DNA variation, human leukocyte antigen (HLA) and killer –cell immunoglobin like receptor (KIR) are remarkable and promising ones (Guha et al., 2013; Moges et al, 2016). Although these markers are immensely supportive, there are few limitations. To cope with this situation, haplotype formed by using autosomal mutations related to the genetic disease is a unique way of studying human migration. The present study reveals the significance of using *cis*-acting mutations of *HBB* gene as the autosomal markers for deciphering human evolution. Moreover, the co-existence of *HBB* mutations can provide valuable information to evaluate the nature and extent of transcontinental admixture in the Indian population.

South Asian population is a rich source of genetic association studies. India is a treasure of population genetics study as different migratory population arrives at different time points and left their genetic imprints (Tamang et al., 2012). Overtime points, intermixing of ANI and ASI occurs in certain cases; and the genetic integrity can not be preserved purely in certain cases. Human culture interacts with genetic variation in a complex way (Creanza and Feldman, 2016). Although, in India, there are social systems, including castes, clans, communities where endogamous marriage is traditionally preferred. Therefore, the efficacy of mutation as well as haplotype study is relevant in the Indian population.

With the gradual progress of various molecular techniques, the field of genetics helps in the development of molecular evolution and anthropology. It is needless to say, further researches and careful investigations are required to decode the full potential of genetic markers in the field of human evolutionary biology. Evolutionary patterns of human migration can be better understood when both genes and culture are considered together. The findings from the present investigation provides valuable baseline data to enlighten the migratory route of the modern human population in India. In future, more genome-wide studies with a larger sample of populations from India will provide further elucidations and intuitions into the population history of India.

## Acknowledgements

The authors are grateful to participants of the present study.

## Consent for publication

All authors have significant contribution to the work and approved the manuscript for submission.

## References

1. Schlebusch CM, Jakobsson M. 2018. Tales of Human Migration, Admixture, and Selection in Africa. Annu Rev Genomics Hum Genet. 19:405–428.

2. Mascie-Taylor CGN, Krzyżanowska M. 2017. Biological aspects of human migration and mobility. Ann Hum Biol 44:427–440.

3. Stringer C. 2002. Modern human origins: progress and prospects. Philos Trans R Soc Lond B Biol Sci. 357:563–579.

4. White TD, Asfaw B, DeGusta D, et al. 2003. Pleistocene Homo sapiens from Middle Awash, Ethiopia. Nature. 423:742–747.

5. Ramachandran S, Deshpande O, Roseman CC, et al. 2005. Support from the relationship of genetic and geographic distance in human populations for a serial founder effect originating in Africa. Proc Natl Acad Sci U S A. 102:15942–15947.

6. Jakobsson M, Scholz SW, Scheet P, et al. 2008. Genotype, haplotype and copy-number variation in worldwide human populations. Nature. 451:998–1003.

7. Jones BL, Raga TO, Liebert A, et al. 2013. Diversity of lactase persistence alleles in Ethiopia: signature of a soft selective sweep. Am J Hum Genet. 93:538–44.

8. Priehodová E, Abdelsawy A, Heyer E, Cerný V. 2014. Lactase persistence variants in Arabia and in the African Arabs. Hum Biol. 86:7–18.

9. Fan S, Hansen ME, Lo Y, Tishkoff SA. 2016. Going global by adapting local: A review of recent human adaptation. Science. 354:54–59.

10. Fernández CI, Wiley AS. 2017. Rethinking the starch digestion hypothesis for AMY1 copy number variation in humans. Am J Phys Anthropol. 163:645–657.

11. Harris K, Pritchard JK. 2017. Rapid evolution of the human mutation spectrum. Elife. 6:e24284.

12. Spichenok O, Budimlija ZM, Mitchell AA, et al. 2011. Prediction of eye and skin color in diverse populations using seven SNPs. Forensic Sci Int Genet. 5:472–478.

13. Nelson MR, Wegmann D, Ehm MG, et al. 2012. An Abundance of Rare Functional Variants in 202 Drug Target Genes Sequenced in 14,002 People. Science. 337:100–104.

14. Reich D, Thangaraj K, Patterson N, et al. 2009. Reconstructing Indian population history. Nature. 461:489–494.

15. Moorjani P, Thangaraj K, Patterson N, et al. 2013. Genetic evidence for recent population mixture in India. Am J Hum Genet. 93:422–438.

16. Basu A, Sarkar-Roy N, Majumder PP. 2016. Genomic reconstruction of the history of extant populations of India reveals five distinct ancestral components and a complex structure. Proc Natl Acad Sci U S A. 113:1594–1599.

17. Mahal DG, Matsoukas IG. 2018. The Geographic Origins of Ethnic Groups in the Indian Subcontinent: Exploring Ancient Footprints with Y-DNA Haplogroups. Front Genet. 23:4.

18. Giardine B, Borg J, Viennas E, et al. 2014. Updates of the HbVar database of human hemoglobin variants and thalassemia mutations. Nucleic Acids Res. 42:D1063–9.

19. Gadhia PK, Vaniawala SN, Kachhadiya TB. 2019. Prevalence of beta thalassemia mutations in population of Gujarat using amplification refractory mutation system-polymerase chain reaction. Int J Community Med Public Health. 6:3294–7.

20. Shah PS, Shah ND, Ray HSP, et al. 2017. Mutation analysis of β-thalassemia in East-Western Indian population: a recent molecular approach. Appl Clin Genet. 10:27–35.

21. Gupta A, Sarwai S, Pathak N, Agarwal S. 2008. Beta-globin gene mutations in India and their linkage to β-haplotypes. Int J Hum Genet. 8:237–241.

22. Nongbri SRL, Verma HK, Lakkakula BVKS, Patra PK. 2017. Presence of atypical beta globin (HBB) gene cluster haplotypes in sickle cell anemia patients of India. Rev Bras Hematol Hemoter. 39:180–182.

23. Sambrook J, Russell DW. 2006. Purification of nucleic acids by extraction with phenol:chloroform. CSH Protoc. 2006:pdb.prot4455.

24. Old JM, Khan SN, Verma I, et al. 2001. A multi-center study in order to further define the molecular basis of beta-thalassemia in Thailand, Pakistan, Sri Lanka, Mauritius, Syria, and India, and to develop a simple molecular diagnostic strategy by amplification refractory mutation system-polymerase chain reaction. Hemoglobin. 25:397–407.

25. Fisher CA, Premawardhena A, de Silva S, et al. 2003. The molecular basis for the thalassaemias in Sri Lanka. Br J Haematol. 121:662–671.

26. Abuzenadah AM, Hussein IM, Damanhouri GA, et al. 2011. Molecular basis of β-thalassemia in the western province of Saudi Arabia: identification of rare β-thalassemia mutations. Hemoglobin. 35:346–357.

27. Patrinos GP, Giardine B, Riemer C, et al. 2004. Improvements in the HbVar database of human hemoglobin variants and thalassemia mutations for population and sequence variation studies. Nucleic Acids Res. 32:D537–D541.

28. Thein SL. 2013. The Molecular Basis of b-Thalassemia. Cold Spring Harb Perspect Med. 3:a011700.

29. Tan JA, Lee PC, Wee YC, et al. 2010. High prevalence of alpha- and beta-thalassemia in the Kadazandusuns in East Malaysia: challenges in providing effective health care for an indigenous group. J Biomed Biotechnol. 2010:706872.

30. Hassan S, Ahmad R, Zakaria Z, et al. 2013. Detection of β-globin Gene Mutations Among β-thalassaemia Carriers and Patients in Malaysia: Application of Multiplex Amplification Refractory Mutation System-Polymerase Chain Reaction. Malays J Med Sci. 20:13–20.

31. Tan JA, Chin SS, Ong GB, et al. 2015. Transfusion-dependent thalassemia in Northern Sarawak: a molecular study to identify different genotypes in the multi-ethnic groups and the importance of genomic sequencing in unstudied populations. Public Health Genomics. 18:60–4.

32. Islam MT, Sarkar SK, Sultana N, et al. 2018. High resolution melting curve analysis targeting the HBB gene mutational hot-spot offers a reliable screening approach for all common as well as most of the rare beta-globin gene mutations in Bangladesh. BMC Genet. 19:1.

33. Gonzalez-Redondo JM, Stoming TA, Kutlar F, et al. 1989. Hb Monroe or alpha 2 beta 230(B12)Arg----Thr, a variant associated with beta-thalassemia due to A G C substitution adjacent to the donor splice site of the first intron. Hemoglobin. 13:67–74.

34. Fedorov AN, Nasyrova FYu, Smirnova EA, et al. 1993. IVS-I-1 (G-->C) in combination with -42 (C-->G) in the promoter region of the beta-globin gene in patients from Tajikistan. Hemoglobin. 17:275–278.

35. Yasmeen H, Toma S, Killeen N, et al. 2016. The molecular characterization of Beta globin gene in thalassemia patients reveals rare and a novel mutations in Pakistani population. Eur J Med Genet. 59:355–362.

36. Hamid M, Shariati G, Saberi A, et al. 2013. Identification of IVS-I (−1) (G > C) or Hb Monroe as a report on the beta-globin gene with a beta-thalassemia minor phenotype in south of Iran. Arch Iran Med. 16:563–564.

37. Fattoum S, Guemira F, Oner C, et al. 1991. Beta-thalassemia, HB S-beta-thalassemia and sickle cell anemia among Tunisians. Hemoglobin. 15:11–21.

38. Win N, Harano T, Harano K, et al. 2002. A wider molecular spectrum of beta-thalassaemia in Myanmar. Br J Haematol. 117:988–992.

39. Ibn Ayub M, Moosa MM, Sarwardi G, et al. 2010. Mutation analysis of the HBB gene in selected Bangladeshi beta-thalassemic individuals: presence of rare mutations. Genet Test Mol Biomarkers. 14:299–302.

40. Moosa MM, Ayub MI, Bashar AE, et al. 2011. Combination of two rare mutations causes β-thalassaemia in a Bangladeshi patient. Genet Mol Biol. 34:406–409.

41. Teh LK, George E, Lai MI, et al. 2014. Molecular basis of transfusion dependent beta-thalassemia major patients in Sabah. J Hum Genet. 59:119–123.

42. Srewaradachpisal K,Tepakhan W, Kanjanaopas S, et al. 2020. Rare β-Globin Gene Mutations including a de novo Mutation of Hemoglobin Hammersmith in Southern Thailand. J Health Sci Med Res. 38:221–229.

43. Gupta A, Hattori Y, Gupta UR, et al. 2003. Molecular genetic testing of beta-thalassemia patients of Indian origin and a novel 8-bp deletion mutation at codons 36/37/38/39. Genet Test. 7:163–168.

44. Meena LP, Kumar K, Singh VK, et al 2013. Study of Mutations in β-Thalassemia Trait among Blood Donors in Eastern Uttar Pradesh. J Clin Diagn Res. 7:1394–1396.

45. Sinha S, Black ML, Agarwal S, et al. 2009. Profiling β-thalassaemia mutations in India at state and regional levels: implications for genetic education, screening and counselling programmes. Hugo J. 3:51–62.

46. Pawar AR, Colah RB, Mohanty D. 1997. A novel beta+-thalassemia mutation (codon 10 GCC --> GCA) and a rare transcriptional mutation (−28A --> G) in Indians. Blood. 89:3888–3889.

47. Colah R, Gorakshakar A, Phanasgaonkar S, et al. 2010. Epidemiology of beta-thalassaemia in Western India: mapping the frequencies and mutations in sub-regions of Maharashtra and Gujarat. Br J Haematol. 149:739–747.

48. Bhukhanvala DS, Italia K, Sawant P, et al. 2013. Molecular characterization of β-thalassemia in four communities in South Gujarat--codon 30 (GL→LA) a predominant mutation in the Kachhiya Patel community. Ann Hematol. 92:1473–1476.

49. Vaz FE, Thakur CB, Banerjee MK, Gangal SG. 2000. Distribution of beta-thalassemia mutations in the Indian population referred to a diagnostic center. Hemoglobin. 24:181–194.

50. Nagar R, Sinha S, Raman R. 2015. Haemoglobinopathies in eastern Indian states: a demographic evaluation. J Community Genet. 6:1–8.

51. Gupta RB, Tiwary RS, Pande PL, et al. 1991. Hemoglobinopathies among the Gond tribal groups of central India; interaction of alpha- and beta-thalassemia with beta chain variants. Hemoglobin.15:441–458.

52. Panigrahi I, Marwaha RK. 2007. Mutational spectrum of thalassemias in India. Indian J Hum Genet.13:36–37.

53. Nishank SS, Ranjit M, Kar SK, Chhotray GP. 2009. Molecular variants and clinical importance of beta-thalassaemia traits found in the state of Orissa, India. Hematology. 14:290–296.

54. Nigam N, Nigam S, Agarwal M, Singh PK. 2017. β-Thalassemia: From Clinical Symptoms to the Management. International Journal of Contemporary Medical Research. 4:77–83.

55. Bakhle A, Dsa L, Bhanushali AA, Nair P, Silveira M, Das BR. 2018. Homozygous hemoglobin Monroe (codon 30 G>C) in a child of Goan (Indian) origin: A case report and family study. J Appl Hematol. 9:151–5.

56. Kulkarni GD, Kulkarni SS, Kadakol GS, et al. 2012. Molecular Basis of β-Thalassemia in Karnataka, India. Genet. Test. Mol. Biomark. 16:138–141.

57. Cordaux R, Stoneking M. 2003. South Asia, the Andamanese, and the genetic evidence for an “early” human dispersal out of Africa. Am J Hum Genet. 72:1586–90.

58. Stanyon R, Sazzini M, Luiselli D. 2009. Timing the first human migration into eastern Asia. J Biol. 8:18.

59. Sharma I, Sharma V, Khan A, et al. 2018. Ancient Human Migrations to and through Jammu Kashmir-India were not of Males Exclusively. Sci Rep. 8:851.

60. Guha P, Srivastava SK, Bhattacharjee S, Chaudhuri TK. 2013. Human migration, diversity and disease association: a convergent role of established and emerging DNA markers. Front Genet. 4:155.

61. Moges AD, Admassu B, Belew D, et al. 2016. Development of Microsatellite Markers and Analysis of Genetic Diversity and Population Structure of Colletotrichum gloeosporioides from Ethipoia. PLoS ONE. 11:e–0151257.

62. Tamang R, Singh L, Thangaraj K. 2012. Complex genetic origin of Indian populations and its implications. J Biosci. 37:911–919.

63. Creanza N, Feldman MW. 2016. Worldwide genetic and cultural change in human evolution. Curr opin Genet Dev.41:85–92.

